# “Primed to Perform:” Dynamic white matter graph communicability may drive metastable network representations of enhanced preparatory cognitive control

**DOI:** 10.1101/2022.09.25.509351

**Authors:** Vivek P. Buch, John M. Bernabei, Grace Ng, Andrew G. Richardson, Ashwin Ramayya, Cameron Brandon, Jennifer Stiso, Danielle S. Bassett, Timothy H. Lucas

## Abstract

Spontaneous neural activity has become increasingly linked to behavioral and cognitive output. A specific cognitive control mode, proactive control, uses prior information to plan and prepare the brain to be particularly sensitive to incoming goal-directed stimuli. Little is known about specific proactive mechanisms implemented via preparatory patterns of spontaneous neural activity, that may enable dynamically enhanced cognitive performance. In this study, humans implanted with intracranial electrodes performed a simple cognitive task. For each subject, pre-trial spectral power and communicability-based features from both grey and white matter nodes were extracted to identify preparatory control states that were “primed to perform”. The anatomical structure and topology of these states across subjects demonstrated a critical role for white matter communicability in decoding and intrinsically controlling preparatory network activity. Our results provide novel insights for putative cognitive network control and may be studied to develop prosthetic approaches for individuals with cognitive deficits.

## Introduction

Spontaneous neural activity, i.e. neural activity not evoked by a specific stimulus or task behavior, has become increasingly linked to behavioral and cognitive output^1–3^. Spontaneous neural activity dynamics have been largely studied using noninvasive neuroimaging modalities in relation to prior task performance. For instance, spontaneous neural activity during rest can encode reactivation of prior task representations for memory formation^4^ and during task behavior can encode immediately prior unconscious errors^5^. Studies investigating pre-stimulus or pre-trial spontaneous neural activity in reference to subsequent activity demonstrate both complex correlates of subsequent stimulus-evoked neural activity^1^ and cognitive behavioral output such as biasing subsequent perceptual inference^6^. In parallel, other studies have shown differences in long time scale spontaneous resting activity between neurotypical and cognitively disabled individuals, such as a decrease in gamma power in individuals with schizophrenia with poor cognitive performance^7^, and alterations in resting state regional activity patterns among Down syndrome patients with cognitive impairment^8^. Further insights into how the brain may be functionally organized to drive subsequent behavior can be gleaned from proactive cognitive control, a major subdivision of cognitive control^9^, whereby neural mechanisms actively maintain relevant information in working memory, trigger goal representations, and coordinate the attention, perception, and action systems in anticipation of subsequent goal-directed activity^10,118^. The mechanisms of proactive control have primarily been studied in the context of task-based conflict, with paradigms such as the AX-CPT^12^, where the proactive period is the time window after an initial stimulus but before the conflict cue is presented. However, proactive control is critical for a wide range of other cognitively demanding events including planning, learning, reasoning, and successful task completion^13,14^; as well as implicated in various neurocognitive and psychiatric disease states^13,14^. It is less known whether or how these proactive mechanisms may be engaged during “preparatory” periods of cognitive activity, just prior to cognitive loading or task engagement, as a behavioral window when specific patterns of spontaneous neural activity may configure brain networks to be particularly ready for upcoming task performance. It is known that resting state activity and functional network architectural profiles can discriminate inter-individual differences in long time scale learning and cognitive output^15,16^. Thus we ask, are there patterns of spontaneous neural activity and network architecture, measured dynamically just prior to cognitive loading, that may drive upcoming cognitive performance? We term this putative interplay between spontaneous neural activity and proactive control mechanisms as preparatory control.

In this study, we therefore aimed to identify within-subject and characterize across subjects, the dynamic, preparatory control states that were particularly “primed to perform”. Using network and spectral analysis techniques on neural activity recorded in 24 patients implanted with stereotactically-placed depth electrodes (SEEG) for epilepsy monitoring, we quantified brain-wide spontaneous nodal activity and graph architecture in the 500 milliseconds prior to task engagement (pre-trial period) as a measure of discrete, potential preparatory periods. We used these discretized brain states to predict performance in the upcoming trial and compared the trial-by-trial preparatory control state to a comparative task-engaged 500 millisecond window just prior to the go cue (intra-trial period). Instead of using a complex conflict discrimination task, we employed a simple temporal expectancy reaction time task (total of 27 task sessions across all patients, with 120-250 trials per session) where the pre-trial period was characterized by a completely blank screen without a fixation cue or any task stimulus, and an intra-trial period in which subjects engaged in a visually-cued instructed delay reaction time task. For each recorded channel (node), we computed spectral power and graph communicability metrics for each frequency band (hereafter referred to as “features”) during the pre-trial and intra-trial periods of each trial, and then identified the nodal features that were dichotomously associated with good (fast) versus poor (slow) reaction time performance in the trial. For each trial, we defined the dynamic preparatory control state space as a concatenated vector of all selected nodal feature values from the pre-trial period that discriminated between good versus poor performance. Preparatory control states that were “primed to perform” were the combination of selected network feature values that preceded good task performance (see Methods). Thus, in our task paradigm, engagement of enhanced preparatory control mechanisms decreased reaction times (RT) and improved task performance, rather than increasing reaction time as seen in conflict-based tasks^17^. Temporal expectancy was utilized for this study due to literature across species identifying neural features predictive of RT during the task-engaged, intra-trial delay period (induced cognitive load)^18–20^. To our knowledge, our study is among the first to examine spontaneous neural features of an internally-driven preparatory period of a temporal expectancy task and compare it to the rich intra-trial period.

Our analyses applied single trial network science techniques to SEEG recordings^21^ to characterize patterns of brain activity and connectivity-based graph architecture. These two types of patterns were integrated to develop a paradigm for detecting and predicting brain states^22,23^. Patterns of brain activity were characterized by computing spectral power (Pow) across the canonical frequency bands^24–27^, while patterns of graph communicability (Qexp) were characterized by first deriving functional connectivity networks in the same canonical bands using phase-locking value (PLV), and then characterizing the architecture of these functional networks using Qexp. There are several graph metrics that describe network and nodal topology (the configuration of a system’s nodes and edges in a network)^28^. We were particularly interested in studying anatomically-derived nodal communication dynamics. Communicability measures the strength of both direct (shortest path) and indirect (non-shortest path) walks between network nodes to assess their capacity for information flow^29,30^ while maintaining the anatomical relationship of these nodes. This differs from latent state space manifolds, where anatomical information becomes hidden^31–34^.

Notably, our characterization of network architecture chose to represent white matter regions as nodes (not edges) within the network. This differs from the commonplace approach of implementing white matter regions as edges in network constructs, since white matter pathways (structural connectivity) constrain functional interactions among brain regions (functional connectivity)^35^. However, since white matter acts as a functional highway for coordinating distributed network processes^36^, and growing evidence also points towards white matter being a source of physiologic and behaviorally relevant functional information^37^, we chose to implement white matter regions as nodes in the network construct with their own activity and architectural profiles. We extracted all potentially robust network features of a preparatory control state space on a within-subject basis, and then comparatively analyzed the topological and anatomical drivers of a metastable state space across subjects. Thus, our study employs a data-driven approach to investigate the potential role of white matter tracts, as well as grey matter regions, in dynamically regulating brain states that are “primed to perform”.

## Results

### Defining the behavioral paradigm and neural activity and architectural metrics

We employed a temporal expectancy task (Figure 1A) in 24 human subjects implanted with intracranial EEG. We defined the pre-trial preparatory period as the 500ms prior to the presentation cue (white box), and a comparison intra-trial task-engaged period as the 500ms prior to the go cue (color change from white box to yellow). We defined network structure by assessing both traditional spectral power-based nodal activity as well as graph communicability, a metric quantifying the potential for frequency-specific information flow through each node of the network. The analyzed functional network structure was therefore assessed by calculating nodal power and Qexp, for each of four canonical frequency bands, averaged in the 500ms pre-trial or intra-trial window (Figure 1B, see methods for detailed description). Each combination of a metric (Qexp or spectral power) and frequency band (theta/alpha, beta, low gamma, high gamma) was defined as a feature. Each recording channel that remained after excluding noisy channels and restricting channels to the ones inside brain tissue on postop imaging was considered individually to be a node, including white matter (see methods). Thus, a node-feature pair (or simply node-feature) refers to a particular feature coming from a specific node.

**Figure 1:**
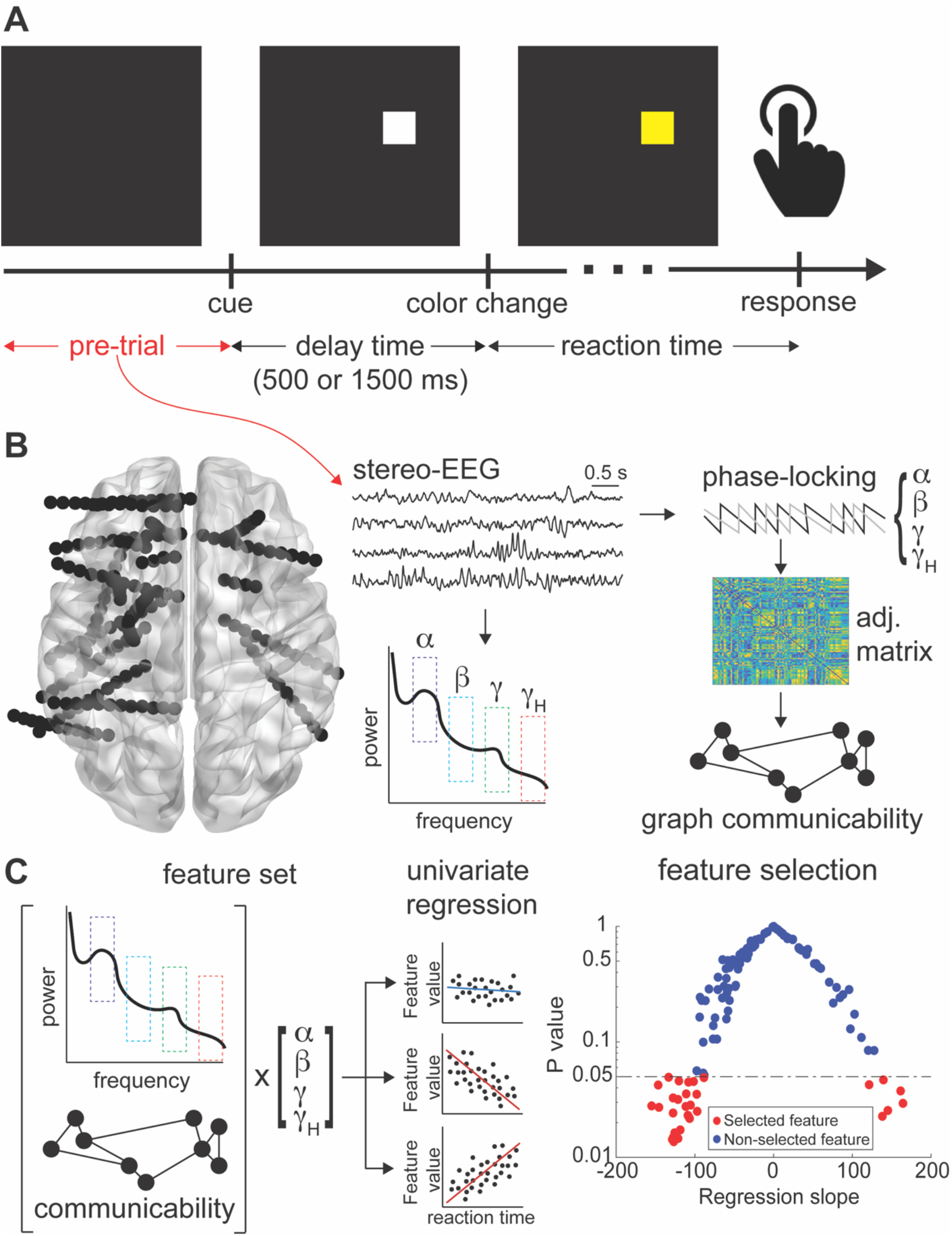
Methods. (A) Temporal expectancy color change detection task. A white cue appears on the screen (presentation cue) in one of nine locations. After a 500ms or 1500ms delay, chosen at random, the white cue changes a yellow cue (go cue). Pre-trial period is defined as the 500ms prior to the presentation cue. Comparative intra-trial period is defined as the 500ms prior to the go cue. (B) Quantifying node-feature pairs as averaged nodal activity (spectral power) or architecture (Qexp) from intracranial EEG during the pre-trial or intra-trial periods. (C) Feature selection. Using an inclusive multiple linear regression approach the complete feature set is reduced to all the node-feature pairs with an uncorrected regression slope p-value of less than 0.05.

### Creating and validating a within-subject cognitive neural state decoder for the preparatory versus task-engaged brain

To identify the specific node-features from each subject that differentiated good versus poor subsequent cognitive performance, thus contributing to a plausible low-dimensional cognitive neural state decoder, we employed a within-subject multiple linear regression feature selection paradigm as a type of demixed dimensionality reduction technique^38^ (Figure 1C). For each subject, brain-wide node-features were calculated in each pre-trial period and regressed against subsequent trial reaction time. Those with univariate regression coefficients that had a p-value of less than 0.05 were initially selected, and concatenated into a vector of preparatory node-features that may contribute to a low-dimensional cognitive neural state decoder for the pre-trial preparatory period (Figure 1C, see methods). This same methodology was repeated for the analyzed network structure during the intra-trial period, where univariate task-engaged node-features were selected and concatenated to represent a plausible low-dimensional task-engaged cognitive neural state decoder.

Next, we tested the capacity for the neural state decoder to predict single-trial cognitive performance by analyzing the z-scored value of each selected node-feature as a trial-by-trial network state space (Figure 2). We demonstrated this approach first in a single subject session. Behaviorally, task performance fluctuated dynamically across trials (Figure 2A). Using only the two most robust node-features to represent the pre-trial preparatory state, a distinction between fast and slow trials emerged (Figure 2B). When all selected node-features were included in the preparatory network state space, it was evident that some node-feature values were elevated prior to fast trials and decreased prior to slow trials, while others had an opposite pattern (Figure 2C). To quantify the single-trial predictive capacity of this state space, a 5-fold cross-validated SVM was trained and bootstrapped over 1000 iterations to assess classification accuracy for ‘fast’ versus ‘slow’ trials (middle third of trials and error trials excluded, see methods). Since selected node-features were those with a univariate relationship to RT, expectedly, classification accuracy in this single subject achieved a robust above-chance performance (AUC = 0.89). Further, a simple null model with the same dataset but randomly shuffling “fast” versus “slow” trial labels led to chance performance (Figure 2D). Using the same methodology, a significant prediction classification was seen in a non-human primate using pre-trial dynamic brain network state structure to predict upcoming trial performance on the same task (Fig S1). Across all subjects, SVM performance was significantly better than chance, and the subject-specific null models expectedly performed at chance (Figure 2E). Validation of the selected network state space SVM was also performed using a temporal partition instead of a 5-fold cross validation approach, beginning with a partition with the first 5% of data in the training set and 95% in the testing set, and ending with a partition with the first 85% of data in the training set and 15% in the testing set. This approach compared SVM performance using the selected network state space, all node features, and the shuffled label null. As expected, with only 5% of training data, the classifier performs near chance, but with increasing amounts of data, the performance asymptotically reaches between 0.7 and 0.8 mean AUC across subjects. Comparatively, using all node features only performs slightly higher than chance, and the shuffled label null performs at chance (Figure S2). To confirm the network state space was more related to upcoming performance rather than previous trial performance, the network state space was also tested as a prediction of the previous trial’s reaction time instead of the upcoming trial. There was above chance performance when classifying the previous trial’s reaction time, but there was high variability and much lower prediction accuracy of the selected network state space for the previous trial RT compared to the upcoming trial RT label (Fig S3). Furthermore, to assess whether the number of selected features in the behavioral dataset was different than what would be expected from randomized data, two additional null model comparisons were tested. Both of these null models were created prior to feature selection, and then the resulting data was passed through the same regression-based machine learning feature selection and network state space extraction pipeline. The first null model was created by randomly shuffling the reaction time values assigned to each trial before selecting features. The second null model was a random alteration to the averaged nodal spectral power and communicability data itself. Multiple univariate linear regression-based feature selection was then performed on these null datasets and duplicated 100 times to create a distribution of features. The number of selected features in the behavioral dataset (n=1452) was significantly higher than the mean number of selected features in the distribution of either null model (one-sample t-test RT randomization t(99))=−36.6, p=2.4 × 10^-59^; Data randomization t(99) = −98.1, p=1.9 × 10^-100^) and in fact lay outside of both distributions entirely (Figure 2E). The RT randomization null distribution had higher variability compared to the data randomization null distribution, though their means were similar (Figure 2E).

**Figure 2:**
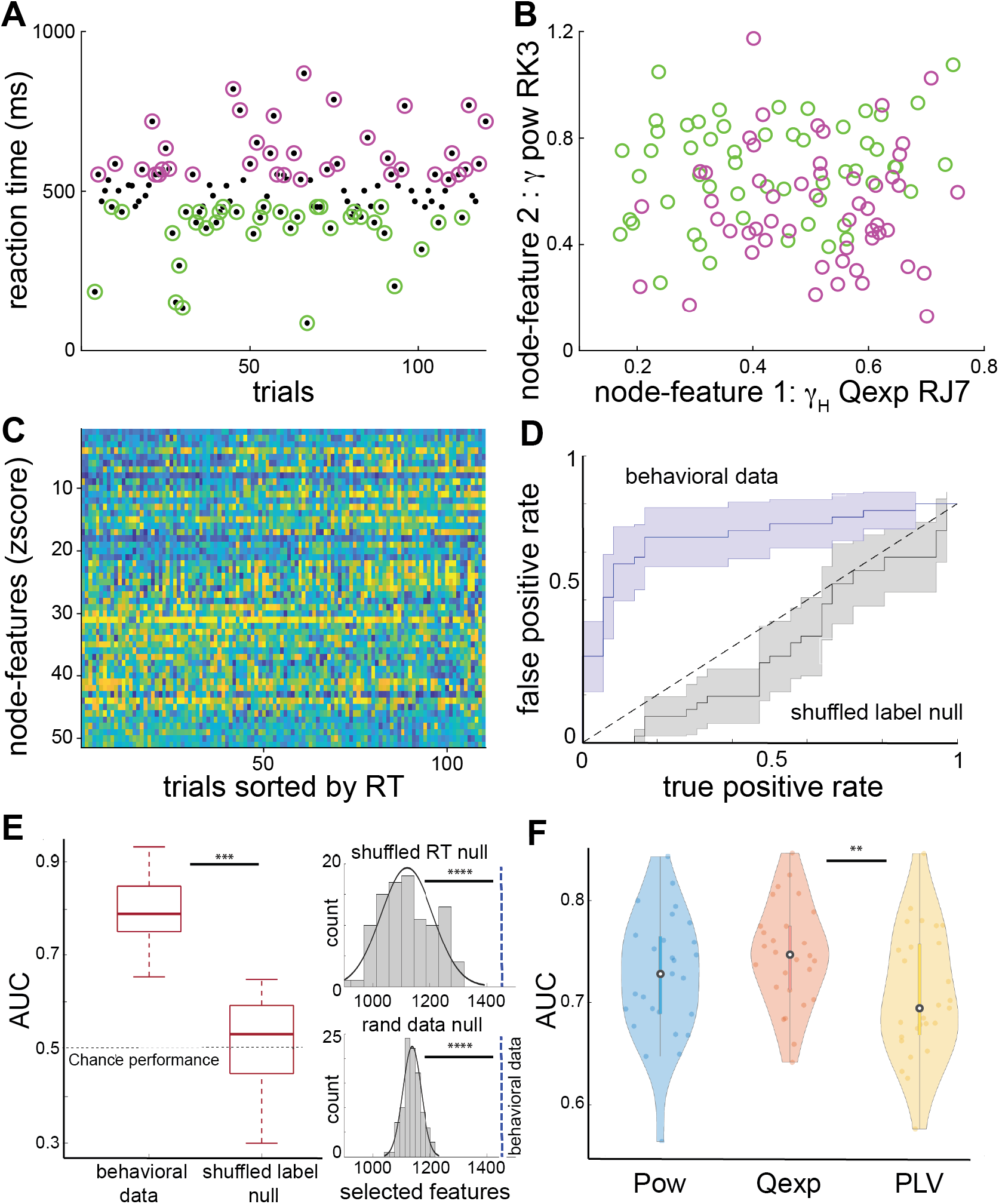
Single-trial brain state classification. (A) Representative patient example of trial-by-trial performance. Magenta designates slowest third of correct trials and green represents fastest third of correct trials. Incorrect trials (RT<0ms or RT>999ms) were excluded. (B) Sample two node-feature space beginning to visually depict RT differentiation between fast and slow trials. (C) Entire extracted node-feature space (within-feature z-scored values across trials to enable direct comparison) plotted against sorted reaction time. X-axis proceeds from slowest to fastest RT trial. Y-axis proceeds from smallest to largest z-score feature value. Some node-features demonstrate low values during slow trials and high values during fast trials (node-features 1-29), while others demonstrate high z-scored value during slow trials and low values during fast trials (node-features 30-51). (D) Single-trial behavioral prediction. Single-trial network state space created as the combination of z-scored values of each extracted node-feature. 5-fold crossvalidated support vector machine (SVM) used to assess trial-by-trial prediction for fastest versus slowest third of trials. Four folds of data randomly chosen to train SVM and one fold used to test prediction. Receiver operating characteristic (ROC) curve demonstrates area under the curve (AUC) performance from 1000 iterations of the SVM test set prediction. Null model created by taking the same data but simply shuffling trial RT label “fast” vs. “slow” randomly. SVM trained on behavioral data performs well in this single subject, while the null model is near chance performance. (E) The distribution of AUC values for the behavioral versus the null dataset across all subjects (n=24). There is a significant difference between the two distributions. Two other types of null datasets were created, this time prior to the multiple univariate regression-based feature selection pipeline. RT randomization assigned RT to each trial randomly prior to single-trial node-feature selection. Data randomization generated a random alteration to the values of each node-feature prior to single trial node-feature selection. Both processes were repeated 100 times to generate a null distribution. Histograms demonstrated the number of selected nodefeatures in both null datasets across 100 iterations across all subjects. Blue line indicates the true number of selected features from the behavioral dataset (n=1452). Using both pre-regression null approaches, the true number of selected node-features in the behavioral data is significantly higher than the null distribution (one-sample t-test RT randomization t(99))=−36.6, p=2.4 × 10^-59^; Data randomization t(99) = −98.1, p=1.9 × 10^-100^). (F) The concatenated features were then segregated into Qexp versus spectral power based features. Additionally, the same regression-based pipeline was performed using nodal phase-locking value instead of Qexp or spectral power as the input data. 5-fold cross-validated SVM was used to compare classifier performance for predicting fast versus slow upcoming trial performance. Across all subjects nodal graph communicability (Qexp) features had significantly higher SVM classifier performance compared to nodal connectivity alone (PLV) (paired Wilcoxon signrank test Zval = 2.81, p=0.0049) and trended towards significantly increased performance compared to nodal power features (paired Wilcoxon signrank test Zval = 1.56, p=0.12).

Next, the subset of selected node features coming from Qexp were separated from the spectral power-based features. A 5-fold cross validated SVM was now trained on each subset of the input data. SVM classification using Qexp nodal architecture as the input trended towards outperforming spectral power-based classification (p=0.12, Figure 2F). The same approach was used to compare classification performance using nodal Qexp as input versus simply using nodal connectivity strength (as measured by PLV) as input. The SVM trained on Qexp features significantly outperformed the SVM trained on PLV features extracted by an identical regression-based approach (p=0.0049, Figure 2F).

### Performance-related differences in preparatory and task-engaged brain network structure

For each subject, we quantified metrics of “ranked Qexp” and “ranked power” by Z-scoring each feature value within-session across all nodes and all trials. The Z-score therefore represents the relative value of a node-feature during a particular trial (Qexp or spectral power in the relevant frequency band for a given node), compared to the mean value of this feature across all nodes and all trials. We thus provide a common dimensional space to plot the ranked Qexp and ranked Pow of each selected node-feature for all subject sessions. Figures 3A, B plot the average Z-score for each selected node-feature across subjects during slow trials against its average Z-score during fast trials. In this plot, color indicated the subject task session (3 subjects performed the task a second time resulting in 27 total sessions across 24 subjects) and size of the marker indicated frequency band of the feature. The y=x diagonal line indicated that a node-feature had the same relative value for both fast and slow trials, and thus is a line of “behavioral equivalence” where fast and slow trials cannot be distinguished. The Euclidean distance from the y=x diagonal was computed as a measure of a node-feature’s average ability to distinguish ‘fast’ from ‘slow’ trials. By comparing the Euclidean distance of ranked node-features derived from pre-trial Qexp compared to power, we found that Qexp node-features had higher Euclidean distances (p<0.0001, Figure 3C), signifying a superior ability to distinguish fast versus slow trials. In contrast, a similar analysis comparing intra-trial Qexp and power node-features found that power node-features had higher Euclidean distances (p< 0.0001, Figure 3C). In other words, Qexp node-features in the pre-trial period better differentiated upcoming performance, while Pow node-features in the intra-trial period better differentiated upcoming performance.

**Figure 3:**
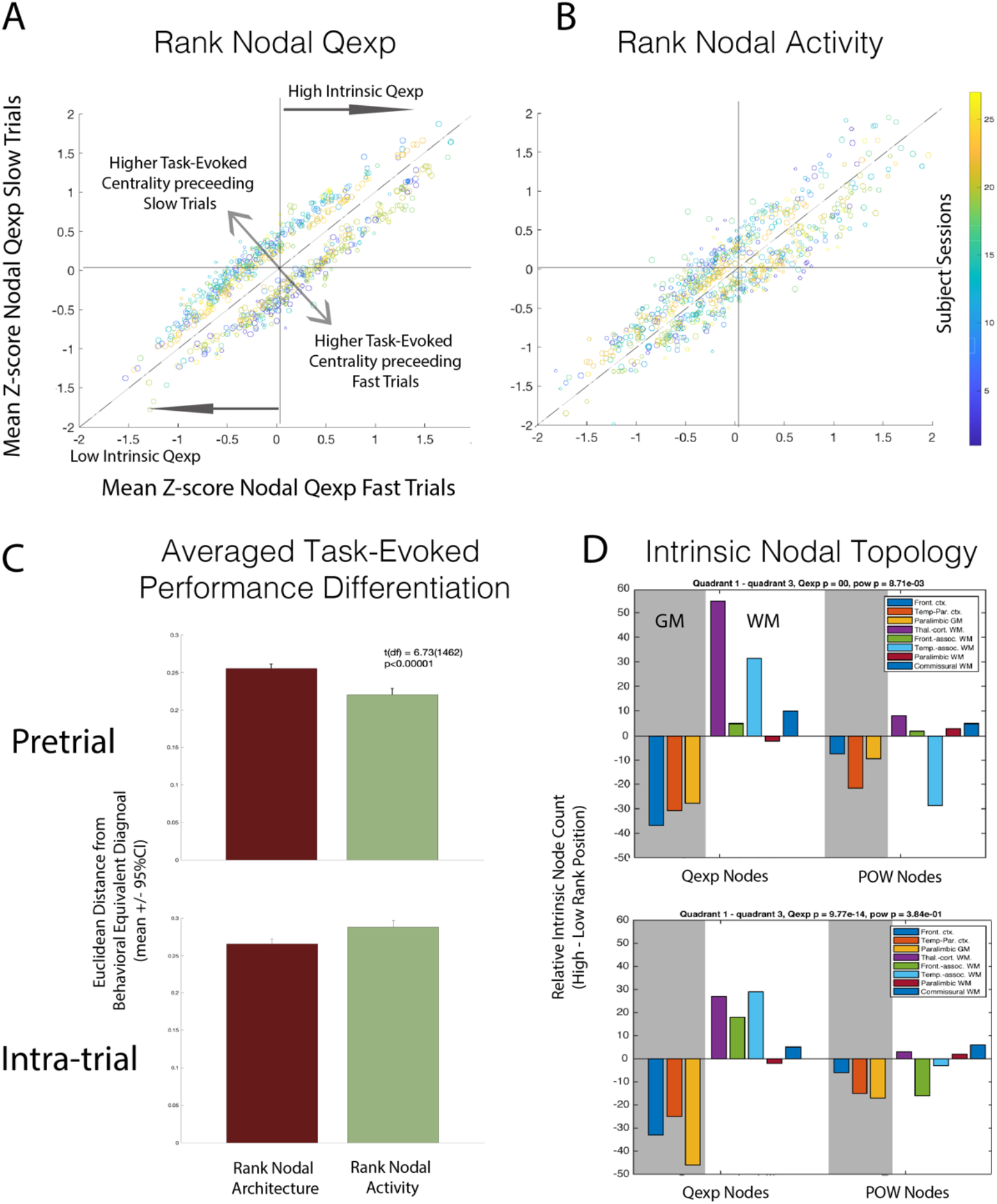
Performance-related and intrinsic network topology. (A) Relationship between ranked nodal commmunicability of extracted Qexp based node-features in the pre-trial period in fast versus slow trials across all subjects. Diagonal line indicates the same z-score value of the node-feature for fast and slow trials. Node-features on the upper side of the diagonal indicate a relatively higher task-evoked value for slow trials. Node-features on the lower side of the diagonal indicate a relatively higher task-evoked value for fast trials. Node-features in the upper right quadrant of the plot indicate that regardless of fast versus slow performance, the feature value is always higher than average (intrinsic high communicability). Node-features in the lower left quadrant indicate the reverse (intrinsic low communicability). (B) Ranked nodal power of extracted power based node-features in the pre-trial period in fast versus slow trials across all subjects. (C) Euclidean distance from behaviorally equivalent diagonal. For the pre-trial period, node-features based on nodal communicability are significantly farther from the behaviorally equivalent diagonal compared to features based on nodal power (p<0.00001). For the intra-trial period, node-features based on nodal power are significantly farther from the behaviorally equivalent diagonal compared to features based on nodal communicability (p<0.0001). (D) Intrinsic nodal topology. During the pre-trial period, node-features based on communicability have high intrinsic communicability in WM, and low intrinsic communicability in GM (chi square test of proportion, statistic = X, p = 7.06e-13.) In particular, thalamocortical WM nodes demonstrate the most robust intrinsic communicability. During the intra-trial period, the pattern still maintains itself between GM and WM nodes; however, the relative difference between the WM pathways (thalamocortical versus frontal and temporal association) becomes more equal. This pattern is not present when defining topology using spectral power based features.

### Intrinsic topology of preparatory and task-engaged dynamic brain network states

We analyzed the intrinsic anatomical topology of the ranked Qexp and ranked Pow node-features by first collapsing the frequency domain, and segregating nodes based on high versus low rank (Figure 3A, B). Node-features in the upper right quadrant have positive z-scored values for both fast and slow trials, suggesting that they intrinsically have high ranked Qexp or power regardless of behavioral performance. Node-features in the lower left quadrant have negative z-scored values for both fast and slow trials, suggesting that they intrinsically low ranked Qexp or power. We asked the question – what was the anatomical distribution of nodes in these two distinct intrinsic topological populations?

To map nodal coordinates to known brain regions, we aggregated grey matter (GM) ROIs from the DKT atlas^39,40^ into three larger regions: (1) frontal cortex, (2) temporoparietal cortex, and (3) paralimbic GM. This atlas was used to segment the brain of each patient. We similarly aggregated white matter (WM) ROIs from the EVE atlas^41^ into five larger regions: (1) thalamocortical, (2) frontal association, (3) temporal association, (4) paralimbic, and (5) commissural WM. The exact ROIs that make up these gross anatomical groupings can be found in Supplemental Table 1. Based on these eight anatomical groupings, we assessed the anatomical distribution of nodes in the upper right versus lower left quadrants of Figure 3A. Surprisingly, we found that nodes with intrinsically high ranked Qexp (upper right quadrant) were predominantly from white matter, whereas nodes with intrinsically low ranked Qexp (lower left quadrant) were predominantly from grey matter (Chi-square test of proportion, p < 0.1e-12). In particular, nodes assigned to thalamocortical white matter had the most robust association with intrinsically high ranked Qexp (Figure 3D). By contrast, for extracted power-based nodes in the upper right and lower left quadrants of Figure 3B, there was no significant difference in the anatomical distribution of nodes with intrinsically high versus low ranked power (Chi-square test of proportion, p = 0.3). In the intra-trial period, the same general pattern remained present, however the ranked nodal Qexp of the thalamocortical WM group was reduced compared to the pre-trial period, and all the WM nodes had similar contribution to intrinsic Qexp (Figure 3D). In other words, the thalamocortical WM nodes had disproportionately high Qexp in the pre-trial period across all trials (fast and slow); while in the intra-trial period nodes across all the major recorded WM pathways (thalamocortical, frontal association, and temporal association) had a similar degree of intrinsically high Qexp.

### Anatomical drivers of the ‘primed to perform’ dynamic brain network state

We then asked if there were any metastable anatomical features predicting single-trial performance across all subjects. We performed a rank-sum test for each feature type (either Qexp or power in each of the four frequency bands, for a total of 8 feature types) and each anatomic region (8 regions total). Thus, we adjusted the significance level to 7.8e-4 to correct for 64 multiple comparisons. With subjects providing repeated observations for the rank-sum tests, we compared the distribution of regression coefficients of all selected node-features within each anatomic region, compared to the distribution of regression coefficients of all non-selected node-features. Thus, this analysis indicated whether a feature type within a given anatomic region can be generalized as a significant positive or negative predictor of reaction time. In Qexp features during the pre-trial period (Figure 4A and B), there was a robust corrected significance for theta/alpha band Qexp in thalamocortical WM nodes as inverse predictors of RT across all subjects (rank-sum p < 1.5e-4), meaning higher values predicted faster RT. Similarly, beta band Qexp measures in paralimbic GM, frontal association WM, and temporal association WM were all significant inverse predictors of RT (rank-sum p < 1.6e-5, 7.8e-4, and 7.8e-4 respectively). Only high-gamma Qexp in frontal GM served as a positive predictor (rank-sum p < 7.8e-4), where higher values predicted slower RT. For power-based features, only nodes in frontal and temporal association WM paths achieved cross-subject significance. High-gamma power in frontal association WM was a significant positive predictor of RT (rank-sum p < 1.5e-4) and beta power in temporal association WM served as an inverse predictor of RT (rank-sum p<1.6e-5).

**Figure 4:**
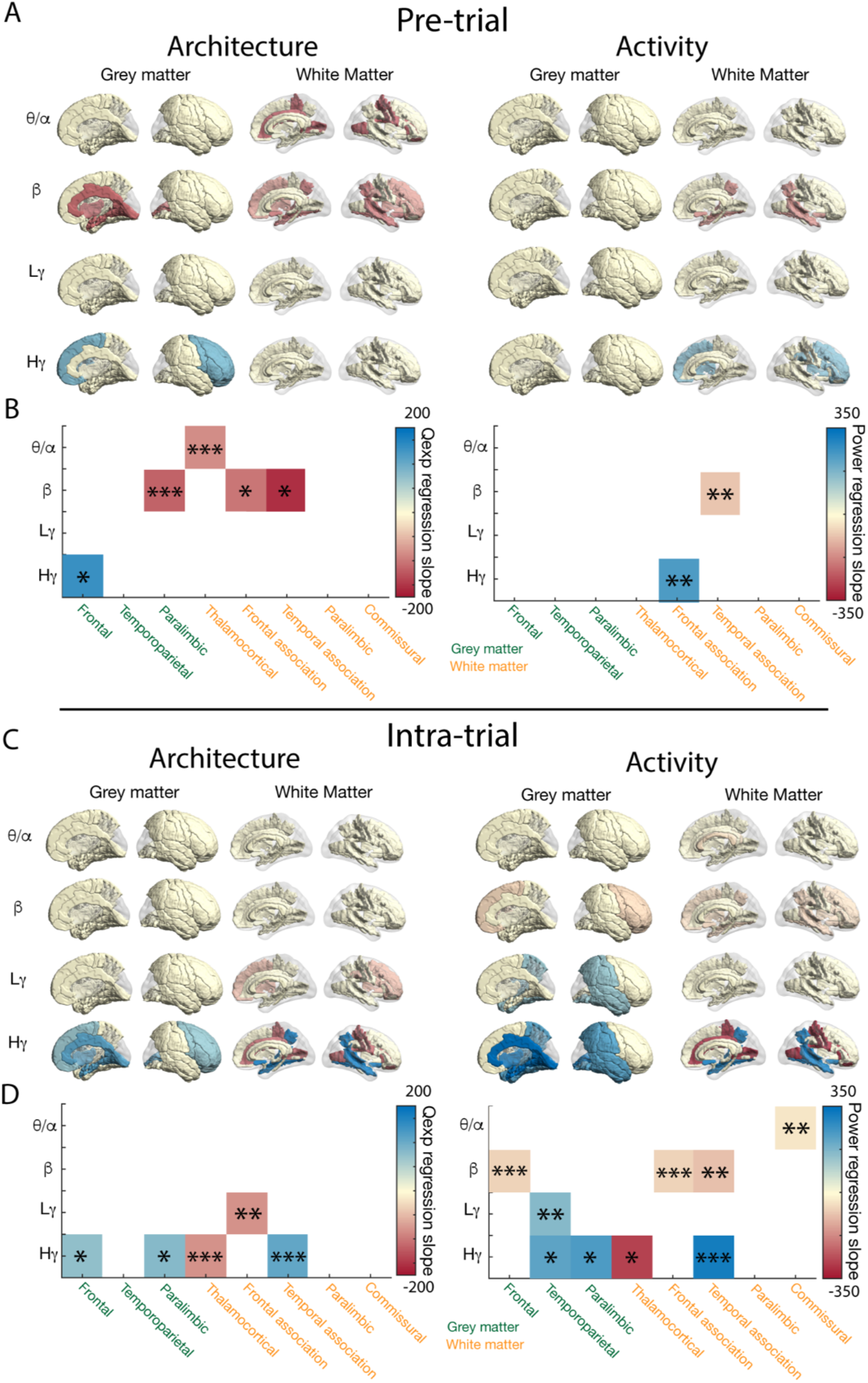
Anatomical drivers of cognitive brain network states across subjects. (A) Pre-trial renderings of anatomical regions from extracted node-features that have a reliable prediction slope for predicting single-trial performance across all subjects (i.e. region-feature pairs; *Left:*nodal architecture, *Right:* nodal activity). Grey matter nodes are designated as being in either Frontal, Temporoparietal, or Paralimbic anatomical group. White matter nodes are designated as being in either Thalamocortical, Frontal association, Temporal association, Paralimbic, or Commissural anatomical group. Red color and intensity indicates degree of inverse relationship with reaction time (larger values associated with faster RT), blue color and intensity indicates degree of positive relationship with reaction time (larger values associated with slower RT). (B) Heatmap display showing all pre-trial region-feature pairs that demonstrate a significant relationship to single-trial performance prediction across all subjects after 64 multiple comparisons correction. Significance testing performed on the distribution of regression slope values for extracted node-features in each region against the distribution of non-selected node-features across subjects. Single asterisk is multiple comparison corrected p<0.05, double asterisk is corrected p<0.01, and triple asterisk is corrected p<0.001. Color designates mean regression slope value for extracted nodes within each region-feature. (C) and (D) Same figures for intra-trial period electrophysiologic data. When compared to the pre-trial period, the anatomical features of single-trial performance prediction from intra-trial period electrophysiologic data convert to being heavily driven by spectral power-based features across grey and white matter regions in beta and gamma ranges. Architecturally, importance is shifted from low frequency to high frequency features in white matter nodes.

Raw data for each feature-anatomy pair in the pre-trial period is found in Supplemental Figure 4. In summary, metastable anatomical drivers of the pre-trial preparatory dynamic brain network state across subjects were primarily from low frequency Qexp, especially in white matter nodes.

As a comparison, we performed the same analysis of region-feature pairs in the intra-trial task-engaged period. The results were largely the opposite of what was seen in the pre-trial period. There was a shift in metastable drivers from low to high frequency, a more even distribution across GM and WM, and predominance of power based features instead of Qexp in the task-engaged dynamic brain network state across subjects. Specifically, for Qexp nodal architecture, we found that frontal association WM in the low-gamma band, and thalamocortical fibers in the high-gamma band were negative predictors of RT (corrected p<0.001 and 0.05 respectively). Meanwhile, frontal GM, paralimbic GM, and temporal association WM were positive predictors of RT in the high-gamma band (corrected p<0.01, 0.05, and 0.01 respectively). However, power was a more broadly predictive metric of reaction time in the task engaged period across all subjects in both GM and WM regions. Theta/alpha power in the commissural fibers, beta power in frontal GM, frontal association WM, and temporal association WM, and high-gamma power in the thalamocortical fibers all served as inverse predictors of RT (corrected p<0.01, 0.001, 0.001, 0.01, and 0.05 respectively). In the low-gamma band, temporoparietal GM served as a positive predictor of RT (corrected p<0.001), while in high-gamma temporoparietal GM, paralimbic GM, and temporoparietal WM served as positive predictors (corrected p<0.05, 0.05, and 0.01 respectively). Raw data for each feature-anatomy pair in the pre-trial period are found in Supplemental Figure S5.

## Discussion

In this study we aim to answer two primary questions: 1) Within each subject is there a specific pattern of preparatory nodal activity and architecture that preferentially enables enhanced dynamic cognitive performance? In other words, for each subject does a dynamic preparatory network structure exist that is “primed to perform?”, and 2) Are there anatomical activity- and/or architecture-based drivers of these dynamic brain network states across subjects? We utilize a simple temporal expectancy task and comparatively analyze predictive dynamic network structure in a 500 millisecond epoch prior to the start of each trial (pre-trial) and prior to the go-cue (intra-trial). By beginning with a within-subject paradigm, followed by an across-subject analysis that explores topological and anatomical properties that are metastable across individuals, we probe the potential network mechanisms underlying preparatory brain states. We perform this analysis using data from the pre-trial versus intra-trial periods to compare the role of preparatory cognitive control during the pre-trial period, versus mechanisms of task-engaged cognitive loading during the intra-trial period. While there are numerous neural signatures from the intra-trial delay period in a temporal expectancy task that are known to predict reaction time^42–48^, less is known about how preparatory brain states (during the pre-trial period) may be optimally organized to be “primed to perform.”

We successfully developed a subject-specific dynamic brain state extraction paradigm that incorporates nodal measures of both neural activity (spectral power) and network architecture (Qexp). Deploying this approach in patients implanted with sEEG allows us to investigate the brain’s preparatory mechanisms for dynamic cognition with high spatiotemporal resolution. When compared to pre-trial PLV (a measure of connectivity strength), we find that pre-trial Qexp architecture provides additional insight into brain communication dynamics, resulting in significantly better performance of a classifier predicting trial-by-trial performance across subjects (Figure 2F). By utilizing a multiple univariate regression approach for feature selection, we specifically create an inclusive but dimensionally reduced nodal feature set with the strongest relationship with reaction time. Since the spatial coverage of channels is heterogeneous between subjects, this type of demixed dimensionality reduction technique efficiently narrows down the number of node-features for each subject according to relevance, while also retaining enough node-features to allow for anatomic overlap across subjects. Further, the regression-based feature selection in particular isolates features that have a dichotomous relationship with RT. Meaning, depending on the sign of the regression slope, a high value of the feature will predict either fast or slow RT, and a low value will predict the opposite. These dichotomously relevant features provide the most robust and efficient differentiation of RT as a within-subject extracted network structure. Therefore, we next ask: what, if any, are the topological and anatomical properties of this extracted network structure that are stable across subjects? And how does this metastable network structure differ between the preparatory pre-trial period compared to the task-engaged intra-trial period?

In the pre-trial period, we find that the relative topological organization of Qexp features across subjects strongly differentiates averaged fast versus slow performance (Figure 3A & C). This implies that relative nodal communicability quantifies aspects of the network’s functional architecture that may predispose subjects to better versus worse performance. By contrast, the relative topology of spectral power-based features are not as robust predictors of performance (Figure 3B, C), implying that nodal activity patterns (as measured by spectral power) are not as effective in differentiating performance states in the pre-trial preparatory period. Interestingly, these findings are completely reversed during the intra-trial period, where nodal activity patterns (instead of architecture) strongly differentiate averaged performance states (Figure 3C).

We next ask the question, are there intrinsic anatomical segregations that contribute to the topological organization of the extracted network structure? We analyze the same dataset of both Qexp-based and spectral power-based node-features, but instead of comparing how these inputs predict fast versus slow performance, we are now concerned with the relative topological organization of these node-features – the landscape of the network where some nodes tend to have higher communicability or power-based activity relative to others, regardless of changes in task behavior. Node-features in the upper right quadrant (quadrant 1) of Figure 3A-B have a positive z-scored value for fast and slow trials. This means that regardless of upcoming trial performance, the relative node-feature value for these particular nodes is always higher than the feature-specific average across all nodes and trials, within each subject. The opposite is true for nodes that fall within the lower left quadrant (quadrant 3). For these nodes, the relative nodal feature value is always lower than the feature-specific average across all nodes and trials, within each subject. This feature specific topology is therefore intrinsic to the node, and not evoked by task performance states. Since z-scores are computed within-subject, the relative topological position in the network is also within-subject and each node’s relative rank can then be concatenated across subjects. This across-subject topological framework can therefore distinguish between nodes that have high or low intrinsic topology in both architectural (communicability) and activity (power)-based feature space. This type of organization, particularly in intrinsic architectural feature space, may inform the controllability of cognitive network structure when introducing specific extrinsic perturbations, such as electrical stimulation, in topologically important nodes^49,50^.

We then collapse our dataset along the frequency domain to focus on identifying any anatomical segregation that may occur with respect to high or low intrinsic nodal position in the feature-specific network topology. Based on post-operative localization, we assign each node to an anatomical group that consists of either frontal, temporoparietal, or paralimbic grey matter versus thalamocortical, frontal association, temporal association, paralimbic, and commissural white matter (see methods). Remarkably, in the pre-trial period there is a near complete dissociation that occurs between low versus high intrinsic communicability based on grey matter versus white matter nodal location (Figure 3D). All grey matter regions have a predominance of Qexp node features in quadrant 3, while nearly all white matter regions have a predominance of Qexp node features in quadrant 1. In particular, thalamocortical projection white matter nodes have the most robust distinction, followed by temporal association white matter nodes. This topological pattern may be present with power-based nodes but is far less robust. Interestingly, temporal association white matter nodes are the only anatomical grouping to have high intrinsic position with architectural features and low intrinsic position with power-based features (Figure 3D). Comparatively, in the intra-trial period, the same overall dissociation remains but now nodes with high intrinsic position are more evenly distributed across thalamocortical projection fibers, frontal, and temporal association white matter regions (Figure 3D). This finding may reflect that the task-engaged brain is facilitating specific network communication and activations requiring involvement of diffuse frontal, temporal, and parietal regions as well as their thalamocortical connections. Further, the fact that there is a robust anatomical distinction between grey and white matter nodes according to their intrinsic network position, particularly with regard to communicability architectural features, may be explained by the fact that white matter acts as a structural bridge connecting large neural populations; thus, white matter likely plays an important role in bridging relevant behavioral networks to construct cognitive brain states.

We then ask the question, do certain architectural or power-based features from specific anatomical regions have a reliable association with trial-by-trial performance across subjects? Surprisingly, we find in the pre-trial period, low frequency communicability in thalamocortical (theta-alpha), frontal association (beta), and temporal association (beta) white matter nodes reliably predicts trial-by-trial performance across subjects (Figure 4A). Specifically, increased low frequency Qexp in these extracted white matter region-features reliably precedes fast performance across subjects (Figure 4A). From this data driven approach, interestingly the grey matter regions that show significant architectural metastable representations happen to be paralimbic grey matter, which includes cingulate gyrus, and frontal grey matter, which includes precentral gyrus, two of the regions thought to have strongest association with proactive control states (Figure 4A). In contrast, the impact of spectral activity-based region features in the pre-trial period is less reliable across subjects (Figure 4A). Interestingly, during the intra-trial period, we find almost the complete opposite (Figure 4B). We find a primary role for spectral activity-based features, as well as a stronger role of the gamma frequency bands in reliably predicting trial-by-trial performance across subjects. Gamma rhythms are known to be associated extensively with perception, attention, and synchronizing local activity^51^, functions which are likely critical for task-dependent cognitive loading during the intra-trial period.

A brain state that is “primed to perform,” in which network communication and neural activity are optimally organized prior to the start of a cognitive task to preferentially enable enhanced performance, relies on the precise interplay of proactive cognitive control mechanisms. The most well-known regions and associated networks involved in proactive control are thought to involve prefrontal cortex, anterior cingulate gyrus, and their respective frontoparietal and cingulo-insulo-opercular networks^52^. These networks are intimately involved with thalamic nuclei particularly in the anterior and mediodorsal nuclear groups^53^. In the pre-trial period, unlike the intra-trial period, we find a primary anatomical driver of the “primed to perform” brain state to be low frequency network architecture in white matter nodes. In particular, these important nodes include thalamocortical white matter pathways known to be linking thalamus to critical hubs in the FPN and CON, as well as frontotemporal associative tracts known to be linking multiple critical cortical regions. Low frequency synchrony is thought to drive inter-regional communication^51^, and theta-alpha range activity has been shown to be particularly important in thalamocortical circuits underlying cognitive functional changes during aging^54^. Interestingly, in our own dataset, the left anterior thalamocortical projection fibers have the most significant contribution to selected node features relative to the sampled distribution across all thalamocortical white matter nodes (Table S2: Left anterior corona radiata percentage of selected features (44/104 = 42.3%) compared to sampled distribution across thalamocortical nodes (117/425 = 27.5%) chi2stat = 7.9350, df=1, p = 0.0048).

These findings have novel implications for the role of white matter in decoding and modulating cognitive states. Although white matter has historically been viewed as neurophysiologically quiet, we find that white matter nodal communicability in the pre-trial period and white matter activity in the intra-trial period are important for predicting task performance. In particular, the finding that thalamocortical WM communicability is the region-feature that is most robust in predicting reaction time across subjects, as well as the finding that thalamocortical WM nodes having the highest intrinsic topology during pre-trial periods, implies that white matter nodes are helpful for decoding cognitive brain states, and potentially also for controlling these brain states with exogenous modulation. These findings are corroborated by literature showing that these may have high structural relevance for cognitively disabled populations; for example, studies in Down Syndrome patients have shown that structural white matter changes in frontoparietal and temporal association and in frontal projection fibers (such as those traveling through the anterior corona radiata) strongly correlate with cognitive disability in these patients^55,56^. In our study, the importance of these white matter nodes likely stems from the tendency of structural connectivity (the anatomical connectome defined by white matter tracts) to shape functional connectivity (the statistical relationships between neurophysiologic events in distinct brain regions) over time^35^. Thus, our findings suggest that certain white matter nodes may be key to identifying brain states, and perhaps also to intervening on brain states.

Our study represents a major step in understanding dynamic mechanisms of preparatory cognitive control, but does have several key limitations. A chief concern is inherent to our method of leveraging SEEG recordings in epilepsy patients. Not only do SEEG implants vary across patients due to unique clinical needs for targeted recordings, but patients with epilepsy frequently have impairments in cognition, as well as distributed abnormalities in both neural activity and network architecture^57^. While these issues affect nearly all studies using human intracranial recordings, they introduce higher sources of within and across-patient variability in our study compared to fMRI studies in healthy subjects. Other limitations include the simplicity of the temporal expectancy task and its study in an unnatural hospital environment, as well as the presence of learning across trial periods, which is known to affect network architecture.

Our study and its limitations prompt several important directions of future research. One direction involves further investigation of white matter electrophysiology as a marker of cognitive state (and predictor of subsequent task performance), and correlating this functional activity with structural connectivity as assessed by personalized diffusion imaging^58^. Second, it may be important to study markers of cognitive control in natural environments by employing intracranial EEG in ambulatory humans outside the hospital. With the availability of chronic implantable neuro-devices (such as the Neuropace RNS and Medtronic RC + S or Percept) that can export data to cloud platforms, such studies are becoming possible^59,60^. Finally, our work opens new directions within the field of brain computer interfaces. Future studies could aim to detect generalized (i.e., not task-specific) “primed to perform” brain states that predict good performance across a variety of cognitive tasks. Additional studies could explore the role of electrical stimulation as a source of network control – that is, exploring whether we can reliably induce network state transitions that push the brain into preferred, “primed to perform” states to improve cognition^61^. These findings would be used to improve early prototypes of closed-loop systems^62^ that detect cognitive states in real-time, *prior* to the start of cognitive loading, and provide targeted stimulation to improve dynamic cognitive output. This type of a closed-loop strategy could have enormous implications for patients with intellectual or cognitive disability, where the overarching goal is not to alter the brain substrate itself, but to preferentially enable functional structures in the substrate that are “primed to perform”.

## Methods

### Subjects and Task

We studied 24 human subjects and one non-human primate with intracranial electrodes that performed the same simple temporal expectancy cognitive task. All human subjects were implanted with stereo-EEG as part of clinical evaluation for epilepsy surgery, and consented to research performed under approval of the Institutional Review Board of the University of Pennsylvania.

Each temporal expectancy trial consisted of a visual cue, a delay period of either 500 or 1500 ms, and a signal after which each subject must press a key in response. We measured reaction time (RT) as the duration between the signal and the time the key was pressed. We eliminated any trials with RT less than zero or greater than one second. Our period of interest for analysis of the ‘pre-trial’ state was the 500ms window prior to the start of each trial during which no stimulus or task-relevant behavior was present. We also analyzed a 500 ms window within each trial, which we refer to as ‘intra-trial’ to validate our approach to mapping activity and architecture, as this period has known anatomic predictors of dynamic cognition.

### Dynamic Brain Network State Modeling

The dynamic brain network state was modeled as the graph architecture and spectral activity of each node (monopolar montage for graph analysis vs. bipolar montage for spectral analysis) across the four canonical frequency bands. Nodes that were outside of the brain and nodes that had significant noise contamination were excluded from each subject’s analysis. For the graph architecture component of network state modeling, to preserve local phase-based activity the raw neural time series data from each contact was processed as a monopolar signal and re-referenced to the common average. A band-pass filter for theta/alpha [3-12Hz], beta [14-30Hz], low gamma [35-55Hz], and high gamma [70-150Hz] followed by a Hilbert transformation was used to calculate frequency-specific instantaneous phase of the raw signal for each node during the time window of interest. Single-trial nodal pairwise phase-locking value (PLV) across this window was then calculated and used for weighted adjacency matrix (W_adj_) construction. We performed this process for both pre-trial and intra-trial epochs.

In order to assess nodal graph architecture incorporating shortest and non-shortest path statistics, pairwise communicability^29,30,63–66^ (Q_exp_) was then calculated by taking the matrix exponential of the PLV-based W_adj_, and the mean Q_exp_ for each node was calculated. For the spectral activity component, a bipolar montage was used to create a time series representing a virtual centroid. Wavelet transformation was used for spectral decomposition and power was computed for each node and then z-scored within the same frequency bands for the same time window as in the graph analysis. Thus, for each graph node (monopolar contact) there was a mean nodal Q_exp_ value for each of the four frequency bands, and for each power node (bipolar centroid), there was a mean spectral power value for each of the four frequency bands. Each nodal metric-frequency combination was considered a feature. We employed a demixed dimensionality reduction technique using linear regression to create a behaviorally-specific low-dimensional state space that extracted and concatenated all the features with a significant (p<0.05) regression slope against RT^67^. This approach enabled a low-dimensional feature space with a preserved relationship between nodal anatomy and the dependent variable of interest, and was used to define each subject’s dynamic brain network state space.

### Quantification of Predictive Performance of Dynamic Brain Network State

The ability for the dynamic brain network state to predict cognitive performance was assessed by splitting subject performance across all non-error trials (RT<0 or RT>999ms were considered error trials) into fastest versus slowest third of RTs. The structure of the dynamic brain network state on a per-trial basis was used to train a support vector machine (SVM). Five-fold cross validation was used to assess performance prediction and this process was bootstrapped 1000 times to generate a receiver operating characteristic curve with 95% confidence intervals. Temporal partitioning instead of 5-fold cross validation was also used with a sliding temporal cutoff between training and test data set to further validate dynamic brain network state performance prediction (supplemental). We trained and tested separate classifiers for pre-trial and intra-trial periods. For a null comparison, the same data was used to train the SVM, but the trial performance assignments (fast versus slow) were randomly shuffled. A second null comparison was also created by using all node features as input into the SVM instead of only using the low-dimensional dynamic brain network state space. Several other null comparisons were tested and are available in the supplemental material.

Next, the subset of extracted node-feature pairs coming from Qexp were separated from the spectral power-based features. A 5-fold cross validated SVM was trained on the subset of extracted Qexp node-feature pairs, and compared to an SVM trained on the subset of extracted spectral power node-features, as well as an SVM trained on a subset of extracted PLV node-features.

### Anatomical localization

Each patient underwent a standard epilepsy imaging protocol including pre-implant MRI, post implant MRI & CT, and post-resection MRI. We used automated warping and labeling (ANTS) to register all images to the pre-implant MRI space^68^. Electrodes were localized and segmented using in-house software^69^, and any electrodes whose centroids fell outside the brain were excluded from analysis.

We also used ANTS to perform a whole-brain automated segmentation using the DKT atlas^70,71^. Localization and segmentations were visually inspected and confirmed for accuracy by a board-certified neuroradiologist. Nodal anatomy was first localized grossly into grey matter versus white matter based on these individual native T1 segmentations. In order to balance increasingly granular spatial resolution with diminishing node counts, ANTS-based anatomical bundles were then created to consist of groups of grey matter and white matter nodes that shared known anatomical or network relationships. We then registered the pre-implant MRI to a ICBM 152 template^72^ in Montreal Neurological Institute (MNI) space and used MNI coordinate to localize structures in white matter using the JHU EVE template^41^ to assign WM electrode contacts to major fiber tracts.

### Anatomical analysis

In order to assess the role of anatomy in the dynamic cognition, we probed whether features in certain regions consistently served to increase or decrease reaction times. To permit sufficient generalizability across patients, we bundled together individual GM and WM ROI to create larger anatomic regions which consist of (i) frontal cortex, (ii) temporoparietal cortex, (iii) paralimbic GM, (iv) thalamocortical fibers, (v) frontal association fibers, (vi) temporal association fibers, (vii) paralimbic WM, and (viii) commissural WM. We list all individual ROI that comprise each composite region in supplemental table 1. Metastable anatomical substates were identified by taking the average regression slope value for each selected node in designated anatomical bundles across all applicable subjects, and testing consistency of direction and magnitude of each anatomical bundle against the null-model of non-selected nodes using the Wilcoxon rank-sum test. We used Bonferroni correction to adjust our significance level for 64 comparisons (8 features × 8 regions).

## Notes

### Competing Interest Statement

The authors have declared no competing interest.

