## Supplemental Materials for "“Primed to Perform:” Dynamic white matter graph communicability may drive metastable network representations of enhanced preparatory cognitive control"

Contents:
Table S1
Table S2
Figure S1
Figure S2
Figure S3
Figure S4
Figure S5

| Composite ROI | Individual ROI |
|---|---|
| Frontal cortex | Superior frontal gyrus, Middle frontal gyrus, Precentral gyrus, Gyrus rectus, Medial orbital gyrus, Posterior orbital gyrus |
| Temporoparietal cortex | Superior temporal gyrus, Middle temporal gyrus, Inferior temporal gyrus, Fusiform gyrus, Angular gyrus, Supramarginal gyrus, Superior parietal lobule, Postcentral gyrus |
| Paralimbic grey matter | Hippocampus, Amygdala, Parahippocampal gyrus, Cingulate gyrus, Lingual gyrus, Entorhinal cortex, Insula |
| Thalamocortical white matter | Anterior/Superior/Posterior corona radiata, Precentral WM, Postcentral WM, Posterior thalamic radiations |
| Frontal association white matter | Superior frontal WM, Middle frontal WM, Inferior frontal WM, Superior longitudinal fasciculus, Superior fronto-occipital fasciculus, External capsule, Lateral fronto-orbital WM, Gyrus rectus WM |
| Temporal association white mater | Middle temporal WM, Inferior temporal WM, Superior temporal WM, Sagittal stratum, Inferior fronto-occipital fasciculus, Fusiform gyrus WM, Angular gyrus WM, Superior parietal WM |
| Paralimbic white matter | Fornix (crus), Stria terminalis, Cingulum (cingulate gyrus) WM, Cingulum (hippocampus) WM, Lingual gyrus WM, Uncinate fasciculus |
| Commissural fibers | Genu/Body/Splenium of corpus callosum |

**Table S1: Anatomic ROI bundles.** We grouped anatomically and functionally linked ROI bundles. The first three regions correspond to grey matter and were determined by a patient-specific segmentation using the DKT atlas. The last 5 regions correspond to white matter and were determined by mapping electrode contacts onto MNI 152 coordinates and using the JHU-EVE template. WM: white matter.

**A)**

| | Number of Selected Nodes | | Number of All Nodes | |
|---|---|---|---|---|
| | *Left* | *Right* | *Left* | *Right* |
| Anterior corona radiata | 44** | 10 | 117 | 53 |
| Superior corona radiata | 4 | 10 | 40 | 36 |
| Posterior corona radiata | 7 | 1 | 11 | 1 |
| Precentral WM | 7 | 13 | 42 | 72 |
| Postcentral WM | 5 | 3 | 26 | 6 |
| Posterior thalamic radiations | 0 | 0 | 15 | 6 |
| **Total** | 104 | | 425 | |

**B)**

| | Percentage of nodes within ROI that are selected | | Percentage of all thalamocortical WM nodes coming from ROI | |
|---|---|---|---|---|
| | *Left* | *Right* | *Left* | *Right* |
| Anterior corona radiata | 37.6 | 18.9 | 27.5 | 12.5 |
| Superior corona radiata | 10.0 | 27.8 | 9.4 | 8.5 |
| Posterior corona radiata | 63.6 | 100.0 | 2.6 | 0.2 |
| Precentral WM | 16.7 | 18.1 | 9.9 | 16.9 |
| Postcentral WM | 19.2 | 50.0 | 6.1 | 1.4 |
| Posterior thalamic radiations | 0.0 | 0.0 | 3.5 | 1.4 |

**Table S2: Breakdown of feature nodes within each thalamocortical WM ROI.** (A) Number of selected nodes and number of all nodes found within each thalamocortical WM ROI (left and right anterior corona radiata, superior corona radiata, posterior corona radiata, precentral white matter, postcentral white matter, and posterior thalamic radiation). (B) Percentage of nodes within each ROI that were selected, and percentage of all thalamocortical WM nodes coming from each ROI.

**Yates corrected chi squared test for proportions: LEFT anterior corona radiata has significantly higher percentage of selected nodes compared to distribution of thalamocortical nodes. 44/104 (42.3%) vs. 117/425 (27.5%) chi2stat = 7.9350, df=1, p = 0.0048. All other nodes were statistically similar to their distribution within the thalamocortical grouping.

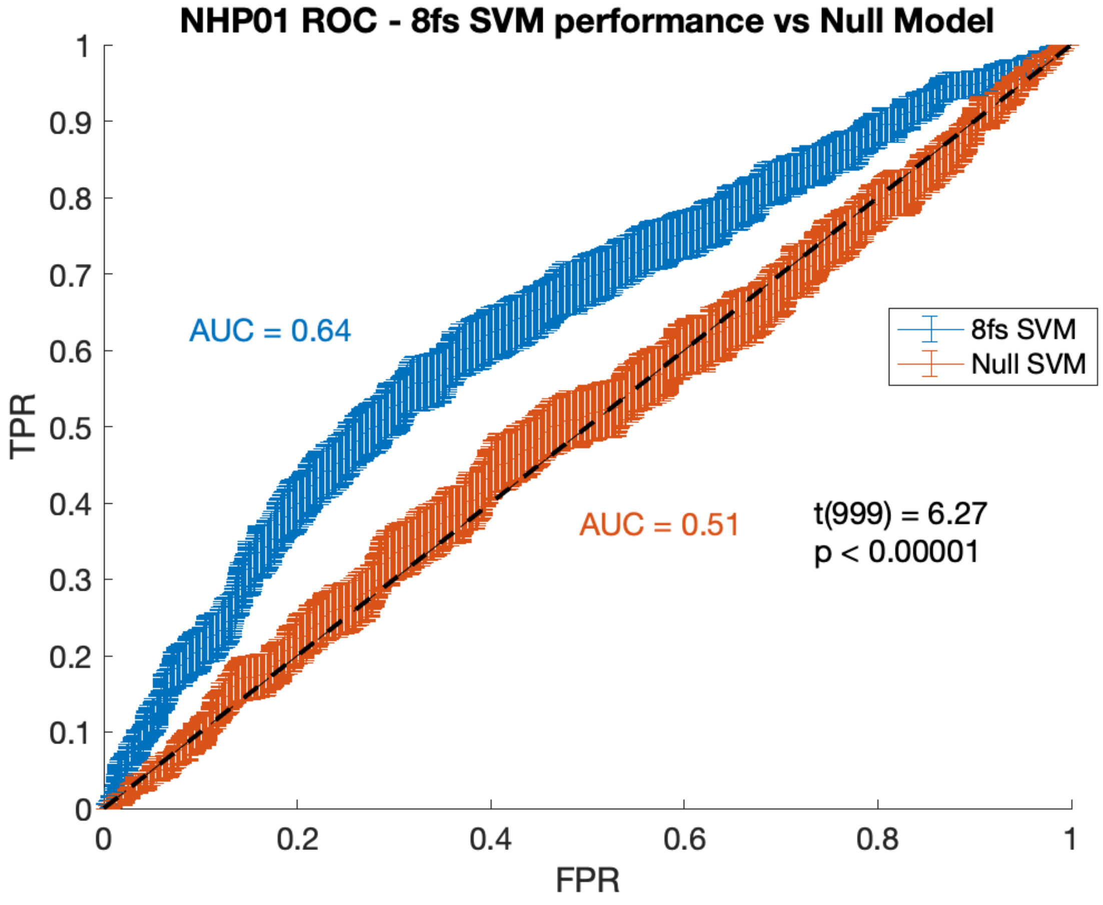


**Supplemental Figure S1: Non-human primate SVM performance.** A support vector machine (SVM) was trained on non-human primate data to perform trial-by-trial prediction of behavior. Input to the SVM was the single-trial network state space, created by combining z-scored values of each extracted node-feature. The SVM performed trial-by-trial prediction of reaction time being in the fastest third versus slowest third of trials, and SVM performance was evaluated with 8-fold cross-validation. The receiver operating characteristic (ROC) curve demonstrates area under the curve (AUC) performance from 1000 bootstrap iterations. For comparison, a null model was created by using the same input data, but with the RT labels being randomly shuffled as “fast” versus “slow”. The SVM trained on behavioral data performs well with an AUC of 0.64, while the null model has near chance performance with an AUC of 0.51.

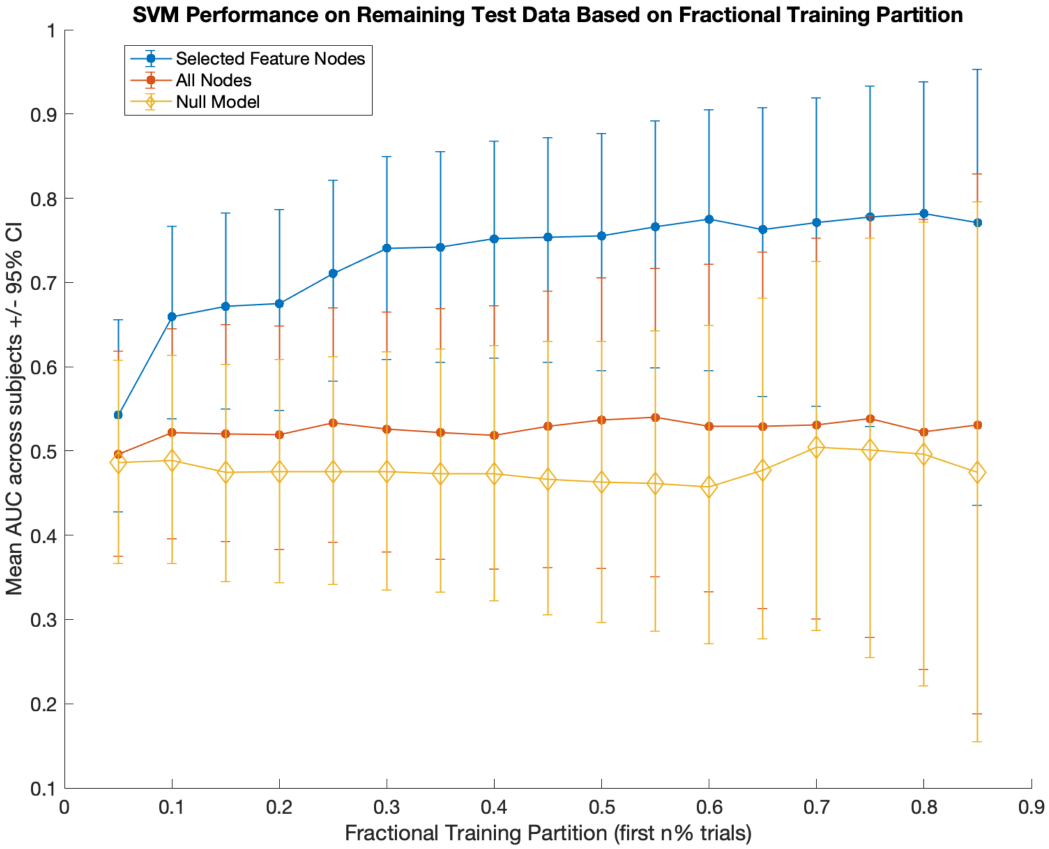


**Supplemental Figure S2: Comparison of SVM performance using data from selected feature nodes, all feature nodes, and null data.** An SVM was trained on progressive amounts of data from human subjects, beginning with a partition with 5% of data in the training set and 95% in the testing set, and ending with a partition with 85% of data in the training set and 15% in the testing set. Mean AUC was averaged across subjects and is plotted with a 95% confidence interval. Blue: performance using selected feature nodes. Red: performance using all nodes. Yellow: performance using null model data.

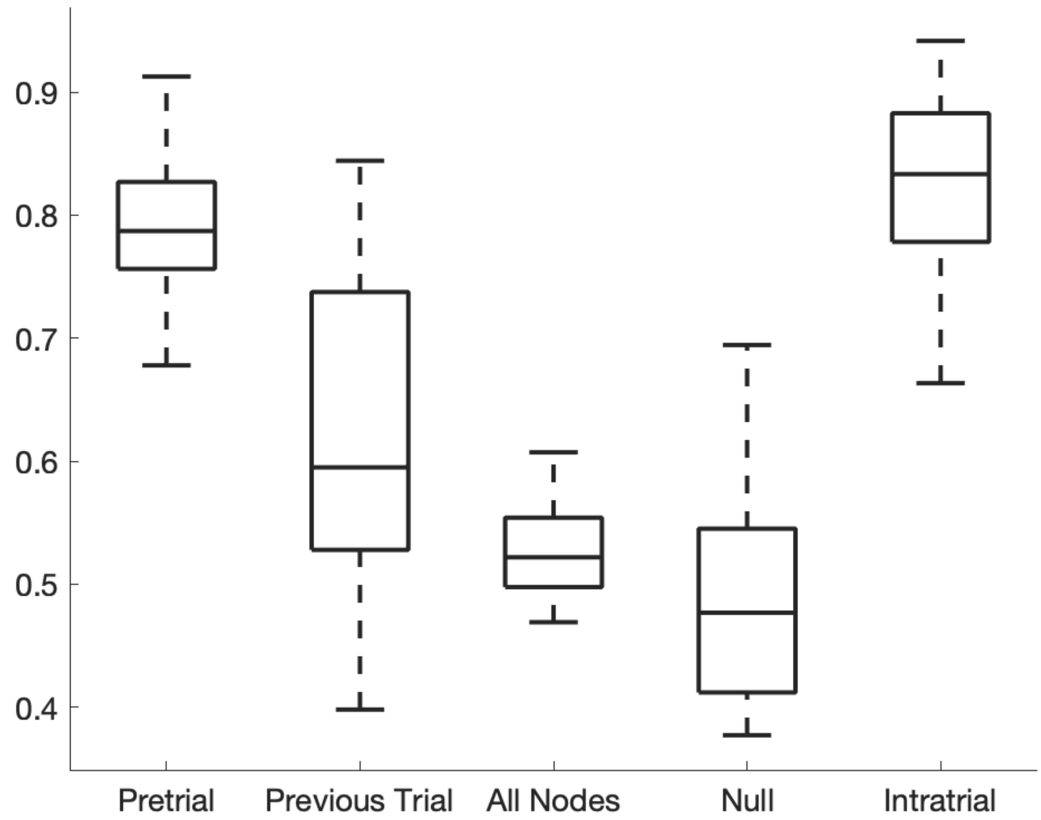


**Supplemental Figure S3: Comparison of SVM performance using various forms of input data.** SVMs were trained to perform trial-by-trial prediction of reaction time being in the fastest third versus slowest third of trials. SVM performance was compared across several types of datasets: data from selected node features during the pre-trial period predicting upcoming trial RT ("Pretrial") as well as predicting the previous trial's RT ("Previous Trial"). Data was also used from all nodes to predict upcoming trial RT ("All Nodes"). The selected node feature data was used with randomly shuffled trial labels of fast versus slow to create a shuffled label null ("Null"). Lastly, selected node features during the intra-trial period were used to predict upcoming RT ("Intratrial").

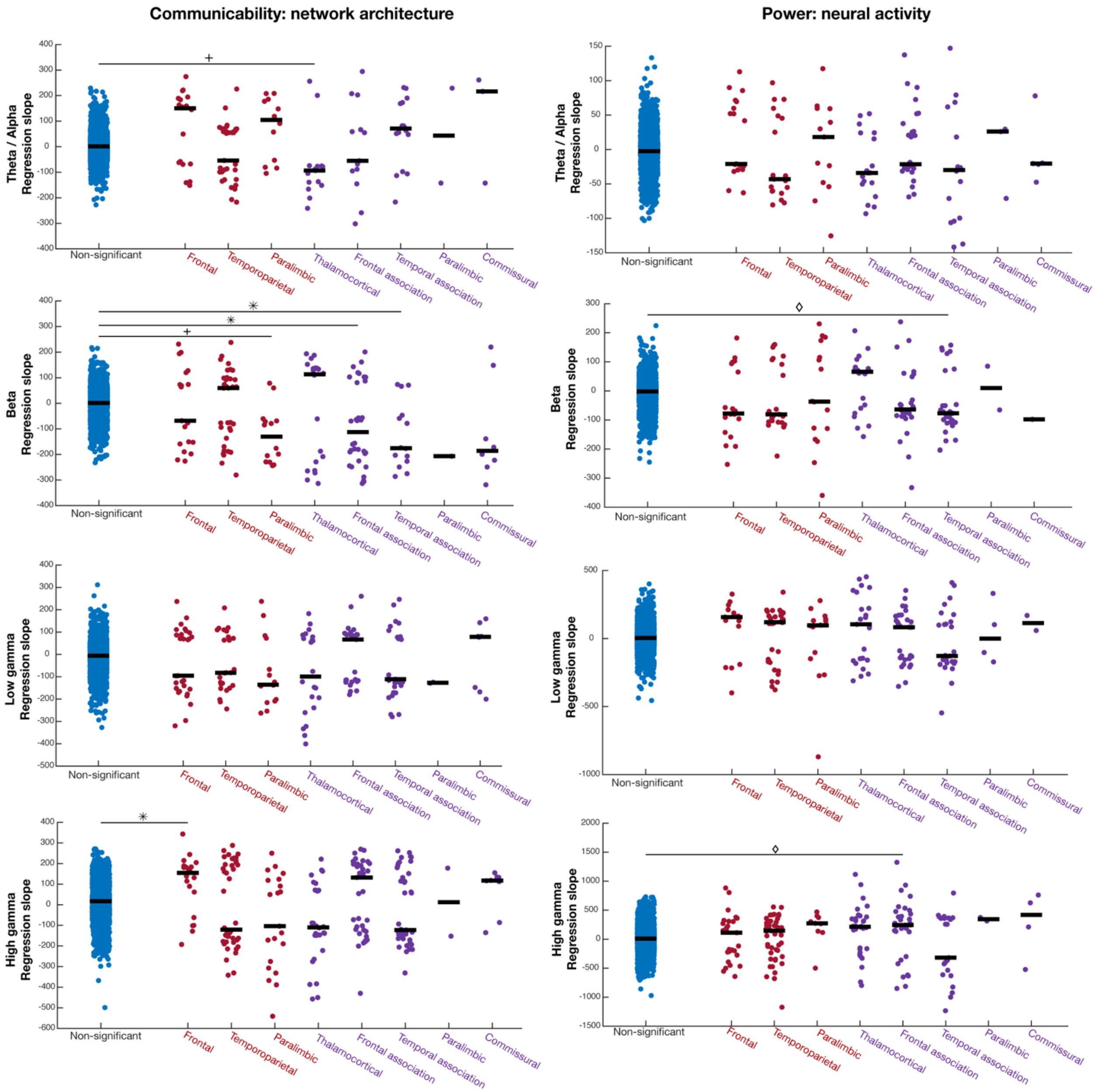


**Figure S4: Metastable anatomic drivers of pre-trial preparatory dynamic brain network state.** Regression slopes of selected nodes (red: grey matter, purple: white matter) in 8 anatomical regions compared to the distribution of non-selected nodes (blue) Left: communicability, Right: power. * = corrected p<0.05, + = corrected p<0.01, diamond = corrected p<0.001.

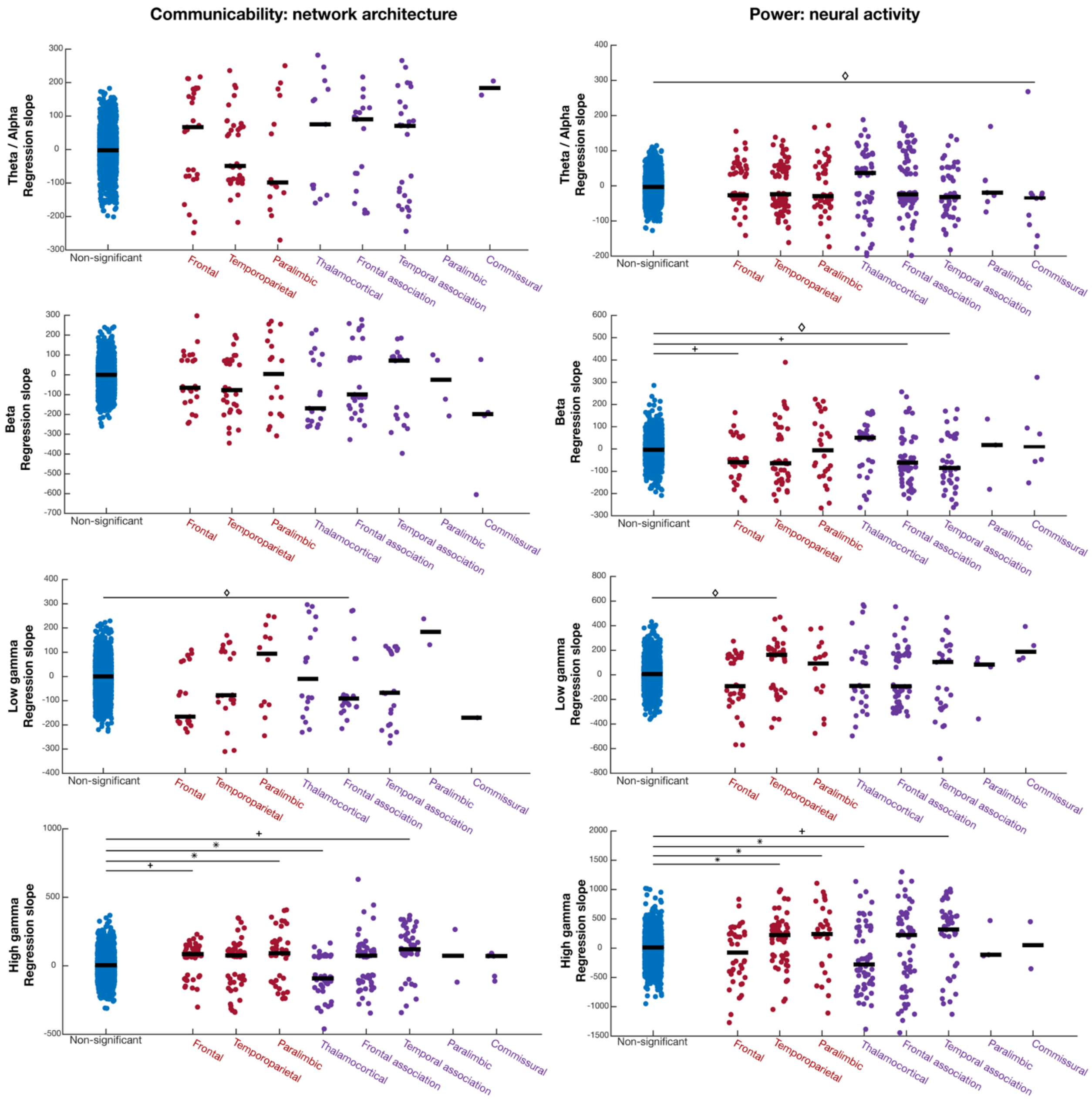


**Figure S5: Metastable anatomic drivers of intra-trial task-engaged dynamic brain network state.** Regression slopes of selected nodes (red: grey matter, purple: white matter) in 8 anatomical regions compared to non-selected nodes (blue) Right: communicability, Left: power. * = corrected $p<0.05$, + = corrected $p<0.01$, diamond = corrected $p<0.001$.
